# AMELIE accelerates Mendelian patient diagnosis directly from the primary literature

**DOI:** 10.1101/171322

**Authors:** Johannes Birgmeier, Maximilian Haeussler, Cole A. Deisseroth, Karthik A. Jagadeesh, Alexander J. Ratner, Harendra Guturu, Aaron M. Wenger, Peter D. Stenson, David N. Cooper, Christopher Ré, Jonathan A. Bernstein, Gill Bejerano

## Abstract

The diagnosis of Mendelian disorders requires labor-intensive literature research. Our software system AMELIE (<u>A</u>utomatic <u>M</u>endelian <u>L</u>iterature <u>E</u>valuation) greatly automates this process. AMELIE parses hundreds of thousands of full text articles to find an underlying diagnosis to explain a patient’s phenotypes given the patient’s exome. AMELIE prioritizes patient candidate genes for their likelihood of causing the patient’s phenotypes. Diagnosis of singleton patients (without relatives’ exomes) is the most time-consuming scenario. AMELIE’s gene ranking method was tested on 215 singleton Mendelian patients with a clinical diagnosis. AMELIE ranked the causal gene among the top 2 in the majority (63%) of cases. Examining AMELIE’s top 10 genes, amounting to 8% of 124 candidate genes with rare functional variants per patient, results in diagnosis for 95% of cases. Strikingly, training only on gene pathogenicity knowledge from 2011 leads to identical performance compared to training on current data. An accompanying analysis web portal has launched at AMELIE.stanford.edu.

## Introduction

Rare diseases, in aggregate, affect 6-8% of the world's population^1^. Patients with Mendelian diseases have one or two genetic mutations in a single gene primarily responsible for their disease and phenotypes^2–5^. Roughly 5,000 rare Mendelian diseases, each with a characteristic set of phenotypes, have been mapped to about 3,500 genes to date^6^. To identify candidate causative genes, exome sequencing is often performed, to relatively high (currently 25-30%) diagnostic yield^7–11^. However, identifying the causal mutations in a patient's exome to arrive at a diagnosis can be very time-consuming, with a typical exome requiring 40 to 100 hours of expert analysis time^12^. This is due to the fact that there are over 4 million variants in a typical human genome by comparison with a reference sequence^13^. Of those, 100-300 are missense or truncating variants that are present, if any, at very low frequency in databases of control individuals^14–16^. These variants are candidates for causing the patient's disease. While in-silico pathogenicity scores^17–22^ keep improving, definitive diagnosis of a known Mendelian disorder is accomplished by matching the patient's genotype and phenotype to previously described cases from the literature. A diagnostic article establishes a causal link between one of the patient's mutated genes and the patient's set of clinical phenotypes.

Because manual literature research is very time-consuming, manually curated databases such as OMIM^23^, OrphaNet^24^, HGMD^25,26^, HPO^27^ and ClinVar^28^ attempt to capture unstructured knowledge from the literature and bring it into a more concise form. The Human Phenotype Ontology (HPO) project^27^ creates two such databases: HPO phenotype ontology, which describes human phenotypes in a structured form, and HPO gene-phenotype annotations (here referred to as HPO-A), containing a list of previously described gene-phenotype relationships curated from OMIM and OrphaNet. Methods such as Phevor^29^, Phenomizer^30^ and others^31–34^ use these databases to prioritize a list of candidate genes for causality given a patient's clinical phenotypes. Dependence on these methods requires continuous comprehensive manual curation, with users ultimately traversing from the ranking tool, through the curating database, in search of the relevant primary literature.

Here, we introduce AMELIE (Automatic Mendelian Literature Evaluation), a method for ranking candidate causal genes directly from the primary literature. AMELIE first automatically analyzes the full text of all relevant papers to create a knowledgebase suitable for diagnosis of patients with suspected Mendelian diseases and then uses this database to automatically prioritize candidate causal genes given the patient's phenotypes. We show that prioritizing candidate causal genes using AMELIE significantly outperforms existing gene ranking methods using a set of 215 clinically diagnosed singleton patients with Mendelian diseases. A web portal for literature based patient analysis has launched at AMELIE.stanford.edu.

## Results

### Knowledgebase construction

We set out to automatically construct a knowledgebase that is suitable for diagnosing patients with Mendelian diseases from the primary literature. To achieve this, we use data from manually curated databases to train a natural language processing system, and apply the system to a comprehensive set of full text articles about Mendelian diseases. An outline of the knowledgebase construction is provided in Figure 1A.

**Figure 1:**
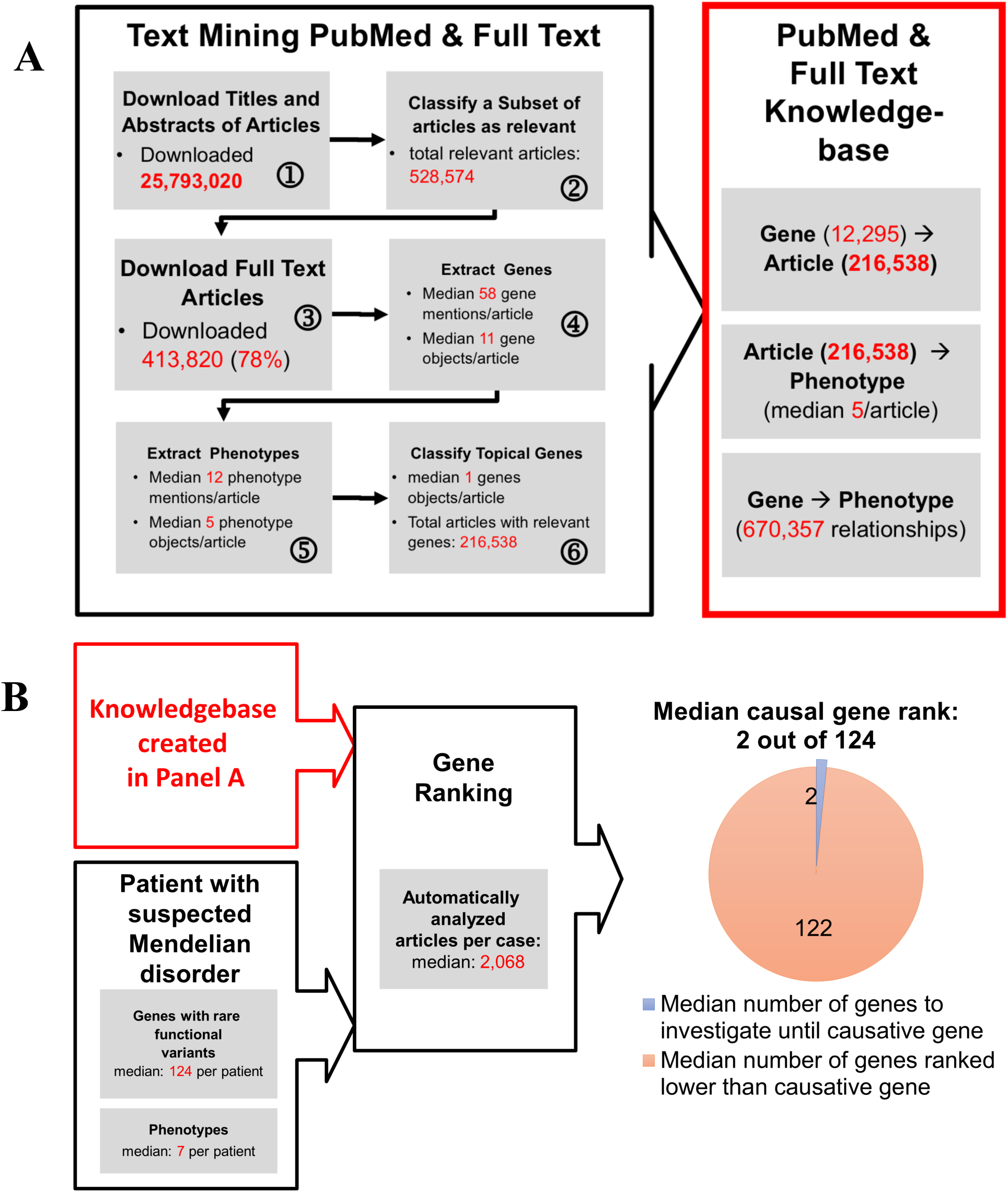
AMELIE (Automatic Mendelian Literature Evaluation) Overview. **(A)** Knowledgebase creation. AMELIE (1) parses all 25.7 million PubMed abstracts, (2) trains a machine learning classifier to detect papers relevant to Mendelian diseases, (3) downloads the full texts of these articles from multiple publishers, identifies (4) gene and (5) phenotype mentions in these texts, and (6) trains a second machine learning classifier to detect which genes may cause which phenotypes, when mutated. The end result (right) is a knowledgebase of 670,357 phenotypes associated with 12,295 distinct genes. **(B)** Candidate gene ranking. AMELIE evaluates each paper in relation to every patient candidate gene with respect to its ability to explain the set of patient phenotypes (see Figure 2). AMELIE reads over 2,000 papers per case, and ranks the causal gene, along with supporting literature, for 215 real patients, as first or second in over 60% of cases (see Figure 3). A median of 122 genes are ranked below the causal gene, greatly accelerating the speed at which a clinician using the system is able to diagnose each case.

#### Downloading titles and abstracts of articles from PubMed (Figure 1A, step 1)

PubMed is a journal citation database of references to peer reviewed biomedical articles containing titles and abstracts of articles. We downloaded all 25,793,020 available titles and abstracts from PubMed.

#### Identifying gene mentions in titles and abstracts

Genes and their protein products are identified by various names in English text. Both the HUGO Gene Nomenclature Committee^35^ (HGNC) and the UniProt^36^ database maintain a list of gene and protein names. To identify mentions of human genes or protein products in the text, a list of 188,975 gene and protein names for 21,346 distinct protein-coding Ensembl genes and 18,149 non-protein-coding genes (which are largely used as negative training examples for the classifiers described below) was compiled from these two sources. AMELIE identifies gene mentions in articles by matching words in the article against the list of gene and protein names and synonyms (Online Methods).

#### Identifying phenotype mentions in titles and abstracts

Human Phenotype Ontology^27^ (HPO) provides a standardized vocabulary of phenotypic abnormalities encountered in human genetic disease. Phenotypic abnormalities are stored with a unique identifier, a canonical name and an optional list of synonyms. Human Phenotype Ontology contains 11,639 distinct human phenotypes. To identify human phenotype mentions in text, a list of names and synonyms of human phenotypes from the Human Phenotype Ontology annotation was assembled comprising 29,182 phenotype names and synonyms. AMELIE identifies phenotype mentions by matching these phenotype names with word groups in English text (Online Methods).

#### Document classification discovers articles about Mendelian diseases (step 2)

PubMed contains titles and abstracts, but not the full text of articles. However, it is the full text that provides the definitive discussion and enumeration of genotypes and phenotypes used to arrive at a paper's conclusions. AMELIE discovers articles relevant to Mendelian diseases using the article's title and abstract, as well as gene and phenotype names discovered in the title and abstract, and then downloads the full text of relevant articles.

We trained a document classifier using logistic regression^37^ featurized by TF-IDF-transformed words (a common transformation of word frequencies into numerical features, after replacing gene and phenotype mentions identified in the titles and abstracts by special tokens (Online Methods). The training set was based on all 51,637 positive articles cited in OMIM's Allelic Variants section and in HGMD and 66,424 random negative articles from PubMed. Articles cited in OMIM and HGMD entries on causative genes for the 215 test patients (discussed below) were omitted from the training data. 5-fold cross-validation of the classifier returned an average precision of 98% and an average recall of 96%. The document classifier was applied to all 25,793,020 titles and available abstracts downloaded from PubMed. Of those, 528,574 (over 10 times the size of the training set) were classified as relevant for Mendelian disease diagnostics.

#### Downloading full text of relevant articles (step 3)

Relevant documents returned from the document classifier were downloaded using PubMunch (Online Methods), resulting in full-text of 413,820 (78%) of all documents classified as relevant. From the full text of these documents, topical genes (which are the gene(s) that are the “topic” of any given article on Mendelian diseases), phenotypes and modes of inheritance were extracted, as described next.

#### Identifying genes and phenotypes in relevant articles’ full text (steps 4-5)

Genes and phenotypes were identified in full text in the same way they were identified in titles and abstracts (above). A median of 58 gene mentions corresponding to 11 distinct Ensembl genes are identified in each article's full text with a precision of 52%. A median of 12 phenotype mentions corresponding to 5 distinct phenotypes are identified in each article's full text with a precision of 92% (Online Methods).

#### Topical gene classification discovers topical genes in text (step 6)

Many gene names mentioned in an article are not relevant for genetic diagnosis. To eliminate these, a “topical gene” classifier was trained to recognize genes that are mentioned in an article as causing a phenotype when mutated. For example, the gene *NOTCH3* is the “topical gene” of the article “Truncating mutations in the last exon of *NOTCH3* cause lateral meningocele syndrome”^38^.

The topical gene classifier is a logistic regression classifier featurized by TF-IDF transformed words flanking all mentions of a gene in an article (Online Methods). The training set was based on 40,350 articles with topical genes that were cited in OMIM and HGMD. All OMIM and HGMD entries on causative genes for the 215 test patients (below) were omitted from the training data. The classifier identifies the topical gene with 91% precision and 83% recall. In total, one, two or more topical genes were extracted from 187,946, 23,308 and 5,284 articles respectively.

#### Linking topical genes to phenotypes

For each article, the topical genes are linked to all the phenotypes mentioned in the article. This way we extracted 670,357 distinct gene-phenotype associations covering 12,295 distinct Ensembl genes.

#### Inheritance mode extraction from articles

AMELIE attempts to extract the inheritance mode from articles. Extracted inheritance modes are either *dominant*, *recessive* or *unknown* (where “recessive” or “dominant” refers to both autosomal and sex-linked inheritance modes). An inheritance mode was extracted from 49,780 (23%) of relevant articles with a topical gene (26,853 with dominant inheritance mode, 22,927 with recessive inheritance mode) at a precision of 98% (Online Methods).

### Patient test set

For evaluation of AMELIE, 215 singleton patients from the Deciphering Developmental Diseases^33^ project were used (Online Methods). The Deciphering Developmental Diseases dataset includes HPO phenotypes (a median of 7 per patient) as well as exome data (VCFs) and the causal genes for each patient (1 per patient). Patient variants were annotated with semantic effect using ANNOVAR^40^ and with frequency information using ExAC^15^ and the 1000 Genomes Project^13^, which combine data from exome and genome sequencing studies from over 60,000 individuals. Rare missense, splice-site, frameshift, nonframeshift indel, stop-gain and stoploss variants were considered to be possibly pathogenic. Genes containing possibly pathogenic variants are considered candidate genes. A median of 124 candidate genes were discovered per patient (Online Methods and Figure 1B).

### Evaluation

As noted above, when training the relevant document classifier and the topical gene classifier described above, all articles about the causal genes in any of the 215 patients were removed from the training data.

#### Phenotype matching calculates similarity of two sets of phenotypes

We associated each phenotypic abnormality in the Human Phenotype Ontology with an information-theoretical score based on the number of genes associated with the phenotype in the knowledgebase we built (Figure 1A). Intuitively, the fewer genes are known to cause a phenotype, the higher the score associated with finding a patient mutated gene capable of explaining this phenotype. Two sets of phenotypes (e.g., one set of patient phenotypes and one set of topical gene phenotypes mentioned in an article) are assigned a *phenotype match score* based on the similarity of the sets of phenotypes and their phenotype scores (Figure 2; Online Methods provide the formal definitions).

**Figure 2:**
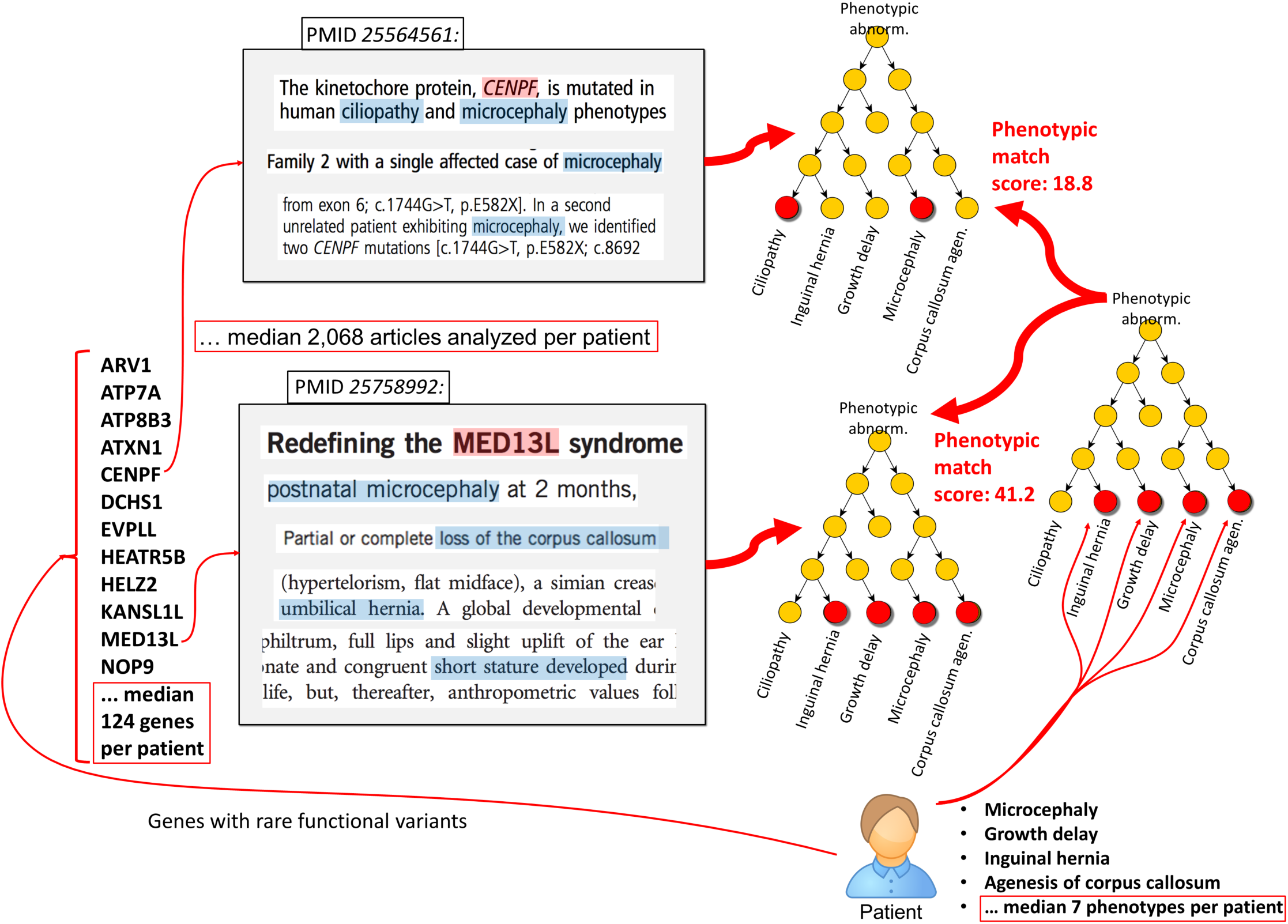
Using machine learning to scan the primary literature in search of a patient candidate gene that best explains the patient set of phenotypes. AMELIE evaluates every paper it has found that ties any of the patient candidate genes to any clinical phenotype/s. AMELIE sums the amount of information acquired by learning from the paper such that the candidate gene can explain one or more of the patient phenotypes (Online Methods). All papers about all candidate genes are scored in this way, and all genes are ranked using their best scoring paper. For each of 215 patients with Mendelian diseases from the Deciphering Developmental Disorders study^39^, AMELIE analyzed a median of 2,068 full text articles about 124 (median) patient genes in an attempt to explain all 7 (median) patient phenotypes.

#### AMELIE ranks candidate causal genes

AMELIE ranks all patient genes for their ability to explain the patient set of phenotypes. Its goal is to rank the causal gene as high as possible, to help minimize the time it then takes the human expert to investigate each of the potentially causal patient variants. Ranking a patient's candidate genes is done from the list of relevant articles annotated with topical genes and phenotypes and possibly an inheritance mode. For each of the patient's candidate genes, AMELIE identifies articles about the gene using the topical gene classifier (above). Each of these articles is matched with the patient's phenotypes. All investigated articles are sorted by their phenotype match score. The patient's candidate genes are then sorted by the highest-ranking article for each gene to provide the reader with an ordered list of possibly diagnostic articles (Figure 2; Online Methods).

### AMELIE outperforms curation dependent methods at ranking candidate causal genes

OMIM^23^ (Online Mendelian Inheritance in Man) is a database of human genes and genetic disorders. OrphaNet^24^ is a database of rare diseases that includes contextual information, such as causal genes and clinical practice guidelines. Both are manually curated from the literature. The set of 215 patients was used to evaluate AMELIE against Phevor^29^ and Phenomizer^30^, both of which use HPO-A gene-phenotype annotations curated from OMIM and OrphaNet as a primary source of information about gene-phenotype links.

For each patient, with a median of 124 candidate genes, AMELIE analyzed a median of 2,068 articles about these genes (Figure 1B). AMELIE ranks the causal gene as the very first gene to read on in 92 out of 215 cases (43%), and in the top 10 genes in 204 out of 215 cases (95%). Phenomizer and Phevor ranked the causal gene at the top in only 72 (33%) and 66 (31%) out of 215 cases, respectively. Similarly, Phenomizer and Phevor ranked the causal gene in the top 10 in only 186 (87%) and 181 (84%) out of 215 cases, respectively (Figure 3). AMELIE performs significantly better than Phenomizer and Phevor (p=8.5*10^−4^ and p=1.7*10^−5^, respectively, one-sided Wilcoxon signed rank test).

**Figure 3:**
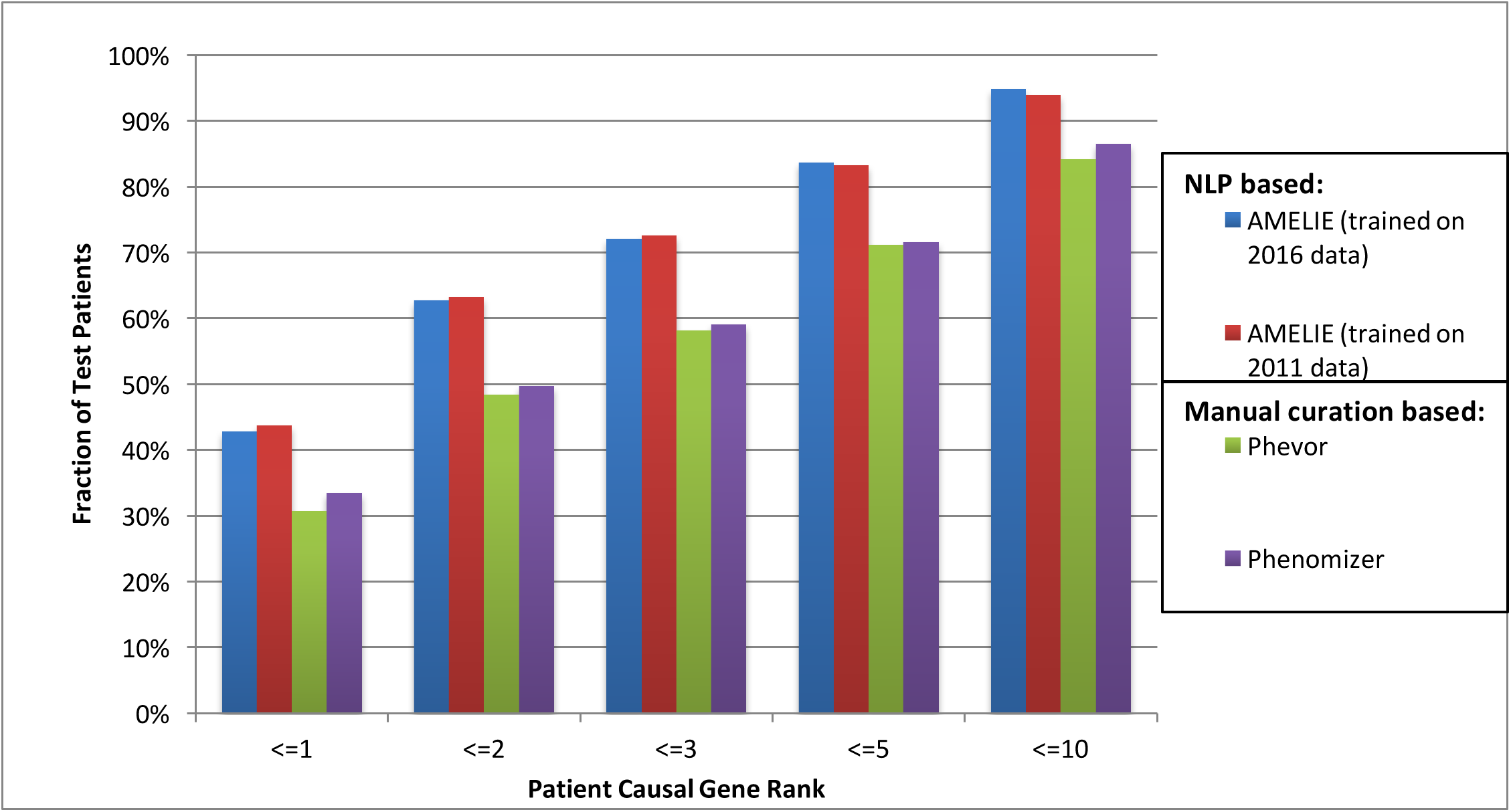
AMELIE patient causal gene ranking outperforms methods based on manually curated databases, even when training only on 2011 knowledge. For each of four methods, the graph shows the fraction of 215 patients with Mendelian diseases, obtained from the Deciphering Developmental Disorders study^39^, where the method places the causal gene in the rank order 1, 1-2, 1-3, 1-5 or 1-10 (left to right). Methods: (Blue, leftmost) AMELIE trained on 2016 data. (Red) AMELIE trained on 2011 data. (Green) Phevor and (Purple, rightmost) Phenomizer, both trained on manually curated OMIM and OrphaNet. Weak supervision using 2011 knowledge (never about the causal genes) is adequate to train AMELIE to be the best performer.

### Performance of AMELIE is unchanged when training only on 5-year-old data

Supervised machine learning methods such as those used by AMELIE rely on a minimum number of labeled training data points to generalize well. Additional data are not needed to improve performance. Here we show that AMELIE generalizes (performs) well already on knowledge available in 2011.

To execute this experiment, the relevant document classifier and the topical gene classifier (above) were both trained only on PubMed titles and abstracts from 2011 or earlier (Online Methods). Impressively, the results on our 215 cases are virtually identical to AMELIE trained on 2016 data (Figure 3).

### AMELIE holds over 3 times as many gene-phenotype relationships as HPO-A

The value of a database that curates gene-phenotype relationships is proportional to the number of gene-phenotype relationships that it contains. Each missing gene-phenotype relationship makes it harder for clinicians and automated patient-solving algorithms alike to correctly associate the causal gene with the patient's disease. HPO-A (build 115) contains an estimated 86,873 true-positive gene-phenotype annotations. Of those, AMELIE recovers 54,371 (68%). In addition, AMELIE extracted an estimated 211,894 true positive gene-phenotype relationships that are not present in the HPO annotations, making it 3.065 times larger than HPO-A (Online Methods).

### Web interface for AMELIE enables patient analysis based on latest literature

A web portal that allows the analysis of patient cases suspected to have a Mendelian disorder utilizing the latest literature has launched at AMELIE.stanford.edu. The portal takes as input a list of genes containing rare variants in the patient and a list of HPO phenotypes observed in the patient. AMELIE article ranking as described above is then performed and a list of genes and top papers is displayed. Clinicians can require certain phenotypes to occur in the article e.g., the clinician could require “hypertrichosis” to show only articles that mention hypertrichosis or any more specific term (child node) in the Human Phenotype Ontology.

## Discussion

We present AMELIE, a method for ranking candidate causal genes and supporting articles in patients with suspected Mendelian disorders. We show that AMELIE ranks the causal gene in the top two in the majority of patients, and within the top 10 genes in 95% of real patient cases. This result suggests that clinicians could rapidly arrive at diagnoses for most diagnosable patients by investigating just the first 10 (8% of the median 124) genes using the relevant literature provided by AMELIE for every incoming patient.

With 5,000 diagnosable Mendelian diseases, caused by roughly 3,500 different genes, and manifesting in different subsets of over 10,000 documented phenotypes, manual patient diagnosis from the primary literature is very labor intensive. Patient phenotypes must be consulted to arrive at a differential diagnosis. One must then read about the different genes known to be involved in these different diseases. Singleton patients may present between 100-300 plausible candidate genes from their exome alone. Memorizing non-obvious phenotype and gene synonyms (e.g., *MLL*=*KMT2A*) makes the task of diagnosis against current literature even harder.

Manually curated databases like OMIM, OrphaNet and HGMD take a step towards alleviating clinician burden by attempting to summarize the current literature. However, manual curation is growing ever more challenging as the literature about Mendelian diseases is increasing at an accelerating rate. Every two years between 2004 and 2015, AMELIE discovered 10% more relevant papers to process. Even more demanding, every single year between 2004 and 2015, AMELIE found nearly 10% more gene-phenotype relationships to curate.

Existing gene- and disease-ranking methods such as Phenomizer and Phevor point clinicians to possibly valuable entries in manually curated databases. To diagnose patients, clinicians will then browse articles linked from these entries in manually curated databases. AMELIE departs from this paradigm. Instead, we propose to automatically pre-process all Mendelian literature to speed up patient diagnosis. By ranking papers directly, AMELIE further accelerates diagnosis. Due to more comprehensive literature analysis (over 2,000 full text articles are analyzed for every patient), AMELIE outperforms these approaches. For example, we manually searched OMIM for the titles of 50 distinct articles that were ranked at the top for the causal genes of 50 random patients. Only 15 out of these 50 articles (30%) are cited in OMIM, as determined by systematic Google searches of the site omim.org on January 30, 2017.

AMELIE's strongest point is perhaps the fact that even when it uses only supervision data from 2011 or earlier, it performs as well as it performs from training on the most current literature. In other words, AMELIE's performance is essentially independent of future manual curation efforts, an important step on the path to offering continuous automated knowledge extraction.

We publish a website at AMELIE.stanford.edu that allows clinicians to interrogate their patients against the latest biomedical literature. Genome-wide data are challenging: no clinician can possibly be expected to memorize the impact of mutations in thousands of different genes. Manual analysis is labor-intensive, slow, costly and irreproducible. Automating as much as possible of this whole process promises to potentiate rapid, affordable, reproducible and accessible clinical genome-wide diagnosis. As such, AMELIE provides an important step on the road to integrating personal genomics into standard clinical practice.

## Acknowledgments

We thank the members of the Bejerano lab, particularly M. J. Berger, H. I. Chen, S. Chinchali, W. Heavner, A. Marcovitz, H. M. Moots, J. H. Notwell, L. Truong, Y. Turakhia, V. Wang and B. Yoo for technical advice and helpful discussions. We thank T. Palomares for assistance with text mining. We thank J. Buckingham and K. MacMillen for assistance with obtaining patient data. We would like to thank the European Genome-Phenome Archive^41^ (EGA) and the Deciphering Developmental Diseases^39^ (DDD) project. The DDD study presents independent research commissioned by the Health Innovation Challenge Fund [grant number HICF-1009-003], a parallel funding partnership between the Wellcome Trust and the Department of Health, and the Wellcome Trust Sanger Institute [grant number WT098051]. The views expressed in this publication are those of the author(s) and not necessarily those of the Wellcome Trust or the Department of Health. The study has UK Research Ethics Committee approval (10/H0305/83, granted by the Cambridge South REC, and GEN/284/12 granted by the Republic of Ireland REC). Deidentified DDD data was obtained through EGA. The research team acknowledges the support of the National Institute for Health Research, through the Comprehensive Clinical Research Network. This work was funded in part by DARPA (CR, GB), the Stanford Pediatrics Department (JAB, GB), a Packard Foundation Fellowship (GB) and a Microsoft Faculty Fellowship (GB).

## Author information

### Contributions

J.B. and G.B. designed the study and analyzed results. J.B. implemented the text mining software, website and associated databases. M.H. authored PubMunch. H.G., A.M.W., J.B., K.A.J., and C.A.D. wrote and improved software tools that were used for genotype and phenotype analysis. C.A.D. analyzed EGA data. A.J.R. wrote parts of the gene and phenotype identification. P.D.S. and D.N.C. curated the HGMD data. C.R. provided text mining guidance. J.A.B. provided clinical interpretation guidance. J.B., M.H. and G.B. wrote the manuscript. All authors commented on and approved the manuscript.

## Online methods

### Knowledgebase construction

#### Downloading titles and abstracts from PubMed (Figure 1A, step 1)

Titles and abstracts from PubMed were obtained using PubMunch (https://github.com/maximilianh/pubMunch).

#### Identifying gene mentions in titles and abstracts

To identify mentions of human genes or protein products in the text, a list of human gene and protein names was assembled using HGNC^35^ symbols, HGNC synonyms, UniProt^36^ gene names and UniProt protein names. Gene mentions are identified in text by matching word groups in the article with gene names from the list of gene and protein names.

To estimate the precision (the fraction of retrieved data points that are true) of the gene identifier, 50 random gene mentions were taken from all downloaded full-text articles and the number of correctly identified genes was counted. A mention was defined as correct if the word group referred to a gene or protein product and the assigned Ensembl gene identifier referred to the mentioned gene.

#### Identifying phenotype mentions in titles and abstracts

To identify human phenotype mentions in the text, a list of names and synonyms of human phenotypes from the Human Phenotype Ontology^27^ annotation build 103 was assembled.

Phenotype mentions are identified by matching word groups in the article with phenotype names from the Human Phenotype Ontology. If a word group matches, it is mapped to the appropriate Human Phenotype Ontology term.

To estimate the precision of the phenotype identifier, 50 random phenotype mentions were taken from all downloaded full-text articles and the number of correctly identified phenotypes was counted. A mention was defined as correct if the word group occurred referred to a phenotype and the HPO ID referred to the mentioned phenotype.

#### Document classification discovers articles about Mendelian diseases (step 2)

To automatically identify articles that are relevant for diagnosing Mendelian diseases, a training set of positive abstracts was created from two existing databases about human mutations. HGMD^42^ (Human Gene Mutation Database) compiles information on mutations that cause human diseases. Its entries contain a gene, the exact mutation, its pathogenicity status and the PubMed ID of the article describing the mutation. OMIM^23^ is a database of Mendelian diseases and genes, as defined in the results section.

The raw text of titles and abstracts was split into sentences and words as described above. Gene and phenotype mentions were identified as described above. All gene mentions in title and abstract were replaced by a token “XGENE” and all phenotype mention in title and abstract were replaced by a token “XPHENO”. All words in the title were prefixed with the characters “TITLE_” and all words in the abstract were prefixed with the words “ABSTRACT_”.

Each document was transformed into a feature vector using the scikit-learn^43^ 0.17.1 CountVectorizer analyzer “word”, an n-gram range of exactly one word and default parameters otherwise. The count-vectorized data was transformed into a TF-IDF feature vector using the scikit-learn^43^ version 0.17.1 TfidfTransformer with default parameters. A scikit-learn^43^ version 0.17.1 Logistic Regression classifier was trained on the TF-IDF feature vector with L2 penalty, a maximum of 1000 iterations and default parameters otherwise.

A TF-IDF transformation treats each document as an unordered bag of words. The document is transformed into a feature vector by assigning each word the scalar product of two statistics: the term frequency (TF) of the word and the inverse document frequency (IDF) of the word. The term frequency *tf(w*, *d)* of a word *w* in a document *d* is defined to be the number of occurrences of *w* in *d.* The inverse document frequency of a word *w* in a document *d* is defined as

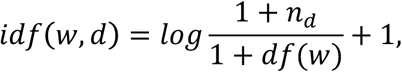

where *n_d_* is the total number of documents and *df(w)* is the number of documents that contain the word *w*. (See also http://scikit-learn.org/stable/modules/feature_extraction.html-text-feature-extraction). Then

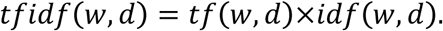

This transformation is applied to each word in a document *d* and inserted in the document's specific feature vector, which is subsequently used by a machine learning model such as the logistic regression models used here.

The document classifier was subsequently run over all titles and abstracts downloaded from PubMed and PubMed IDs for relevant articles were returned.

#### Downloading full text of relevant articles (step 3)

Relevant documents returned from the document classifier were downloaded using PubMunch. Downloaded articles in PDF format were converted to text using pdftotext version 0.26.5 (https://poppler.freedesktop.org/). From the full text of these documents, topical genes, phenotypes and disease inheritance modes were extracted.

#### Identifying genes and phenotypes in relevant articles’ full text (steps 4-5)

Genes and phenotypes were identified in full text in the same way they were identified in titles and abstracts as described above.

#### Topical gene classification discovers topical genes in text (step 6)

To identify which genes are causing human phenotypes when mutated, a “topical gene” classifier was trained. The topical gene classifier takes as input all gene mentions from the article's full text (which are discovered as described above) and identifies which gene object mentioned in the article is causing phenotypes when mutated.

To construct the training set, for articles cited in the OMIM Allelic Variants sections, the topical gene in each article is the OMIM gene entry from which the article is cited. For articles cited in HGMD, the topical gene is the gene deposited in HGMD. All OMIM and HGMD entries on causative genes for the 215 test patients were omitted from the labeled data set. The labeled data set contains 43,228 positive examples of topical genes from 40,350 articles. The negative set consists of all gene objects mentioned in an article that are not the topical gene and consists of 569,747 negative training examples. The whole training set was split into 28,262 (roughly 70%) articles used for training (“training set”), 4,017 (10%) articles for running evaluation and improvement of the classifier (“development set”), and 8,071 (20%) articles for final performance testing (“test set”).

The classifier is then trained on the following features:

- Number of mentions of the gene in the title
- Number of mentions of the gene in the abstract
- Number of mentions of the gene in the full text
- TF-IDF-transformed word counts (defined above) in 5-word-windows around all gene mentions for the gene in question

A scikit-learn^43^ 0.17.1 logistic regression classifier with default parameters is subsequently trained on these features, and the classifier is applied to all gene objects in all downloaded articles and all PubMed titles and abstracts. Topical gene mentions from PubMed title/abstracts and full text are subsequently combined for each relevant article. Articles in which more than 10 topical genes were identified were omitted from the knowledge base. Precision (the fraction of retrieved data points that are true) and recall (the fraction of all true data points that were retrieved) of the classifier were determined by running the classifier on the test set.

#### Linking topical genes to phenotypes

AMELIE gene-phenotype extractions were compared with gene-phenotype extractions curated by HPO build 115 (the latest build available in February 2017), after canonicalization (see forth) of both sets. Phenotypic abnormalities in HPO are structured as a directed acyclic graph (DAG) with a single root. “Canonicalization” of a set of phenotypic abnormalities in the HPO ontology refers to the following process: a set of phenotypic abnormalities (i.e., descendants of the node “Phenotypic Abnormality”, HP:0000118) is augmented by adding all its ancestors up to and including “Phenotypic Abnormality”. A canonicalized set of gene-phenotype relationships refers to a set of gene-phenotype relationships where each gene-phenotype link is augmented by gene-phenotype links for all ancestors of the phenotype up to “Phenotypic Abnormality”. E.g., if the original set of gene-phenotype links includes “*KMT2A* – Elbow Hypertrichosis”, then the canonicalized set of gene-phenotype links includes “*KMT2A* – Hypertrichosis” etc. up to “*KMT2A* – Phenotypic Abnormality”.

#### Inheritance mode extraction from articles

For each article, AMELIE attempts to extract mentioned inheritance mode(s). If the title and abstract of an article contain only words indicating that the described disease(s) are inherited in a *dominant* fashion (“heterozygous”, “heterozygote”, “heterozygosity”, “dominant”, “dominantly”, “autosomal-dominant”), then the article is assumed to describe a dominantly inherited disease. If the title and abstract of an article contain only words indicating that the described disease(s) are inherited in a *recessive* fashion (“homozygous”, “homozygote”, “homozygosity”, “recessive”, “recessively”, “heteroallelic”, “autosomal-recessive”, “biallelic”, “compound heterozygous”), then the article is assumed to describe a recessively inherited disease. If the inheritance mode cannot be uniquely identified from title and abstract using this method, the inheritance mode described in the article is extracted as *unknown*.

An analysis of 50 random articles about Mendelian diseases with an extracted inheritance mode revealed that 49 out of 50 were assigned correctly (precision: 98%).

### Patient test set

VCF files of patients submitted to the Deciphering Developmental Disorders^39,44,45^ (DDD) project were downloaded from the European Genome-Phenome Archive^46^ (EGA). The EGA accession numbers were EGAD00001001848, EGAD00001001977, EGAD00001002748, EGAD00001001355, EGAD00001001413 and EGAD00001001114. All patients with a single-gene diagnosis, also found in their VCF, that was not due to a structural variant and for which the causative gene was not a novel discovery of the DDD project were selected. From any diagnosed set of siblings, a single diagnosed sibling was selected at random. This resulted in an intermediate set of 223 diagnoses.

#### Variant filtering defines the candidate gene list for each patient

ANNOVAR^40^ v527 was used to annotate variants with predicted effect on protein coding genes using gene isoforms from the Ensembl gene set version 75 for the hg19/GRCh37 assembly of the human genome^40^. All coding isoforms were used where the transcript start and end are marked as complete and the coding span is a multiple of three. Patient variants are annotated with frequency information from ExAC^15^ and the 1000 genomes project^47^, as previously described in ^48^.

To obtain a candidate gene list per patient, variants fulfilling the following criteria are assumed to be possibly pathogenic: (a) The variant is in the coding region of a gene or in a canonical splice site and not synonymous. (b) The overall allele frequency in both the ExAC and the 1000 Genomes control populations does not exceed 0.5% and the homozygote count is not greater than 1. (c) Variant calls with inconsistent ALT variant calls (2 or more lines in the same VCF with different alternative allele calls) and variant calls with inconsistent REF calls (2 or more lines in the same VCF with different reference allele calls) are removed. (d) For transcripts with a single heterozygous variant, the frequency of the variant in ExAC and the 1000 Genomes Project has to be 0.1% or less and the allele count has to be 3 or less. Using this filtering scheme, 8/223 (3.5%) of diagnoses were flagged where the reported causal variant/s occur/s in a significant number of presumably non-affected individuals in ExAC. The final test set we used consists of the remaining 215 patients.

### Evaluation

#### Phenotype matching calculates similarity of two sets of phenotypes

Each phenotype node *x* in the Human Phenotype Ontology that is a descendant of HP:0000118 (“Phenotypic Abnormality”) is associated with an information-theoretic score. The score of *x* is calculated as

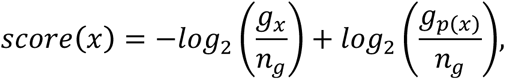

where *g_x_* is the number of genes we have learned can cause *x*, *n_g_* is the number of genes we have learned can cause any HPO phenotype and *g_p(x)_* is the number of genes we have learned can cause all parents of *x* in the HPO directed acyclic graph (DAG). By virtue of the HPO DAG structure design, when a gene causes some phenotype *x*, it also causes all ancestors of *x* up to Phenotypic Abnormality. E.g., if mutations in *KMT2A* cause “elbow hypertrichosis”, then mutations in KMT2A also cause “hypertrichosis”, “abnormal hair quantity”, “abnormality of the hair”, “abnormality of skin adnexa morphology”, “abnormality of the integument”, and the most general term, “phenotypic abnormality”.

To calculate the match score between two sets of phenotypes *A* and *B* (e.g., the match score between a patient's set of phenotypes *A* and an article's set of phenotypes B), let *A*’ = *A* + *all ancestors of nodes in A* and similarly *B*’ = *B* + *all ancestors of nodes in B*. Let *C* = *intersection of A’ and B’*.

The match score of the sets of phenotypes *A* and *B* is defined as

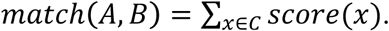

#### AMELIE ranks candidate causal genes

The output of the automated ranking system is the list of patient mutated genes, ranked by their likelihood of individually causing the case's phenotype. To generate this gene list, all articles about one of the patient's genes carrying rare non-synonymous variants are examined. An article *A* is assumed to be “about” a gene *G* if the gene *G* is identified as a “topical gene” from article *A* by the topical gene classifier (above). If a gene contains a single heterozygous variant, but the article describes a recessive disease, the article is omitted from the analysis. Each of the examined articles receives a phenotype match score that is calculated by matching all the phenotypes associated with the topical gene in the article with the case's phenotypes using the *match* formula described above.

The output of the solver is a list of genes associated with articles, sorted by the phenotype match score of the highest ranked article for each gene. In rare cases, multiple articles (for the same or different genes) receive equal match scores. To break tied match scores, additional sorting criteria are applied:

The RVIS^49^ score is a measure of a gene's intolerance to nonsynonymous variants derived from population frequencies of synonymous and nonsynonymous variants in a gene. Genes with low RVIS scores are likely to be intolerant to nonsynonymous variants. Genes with high RVIS score are more tolerant to such variants.

To break rare ties between articles’ phenotypic match scores, the following additional sorting criteria are applied: (2) the RVIS score of the mutated gene (lower RVIS scores are ranked higher) (3) the publication year and month of the article (newer articles are ranked higher) and (4) by the unique PubMed ID of the article.

### AMELIE outperforms curation dependent methods at ranking candidate causal genes

#### Comparison to Phevor

The output of Phevor^29^ Version 2 for each of the 215 patients was obtained through Phevor's website (http://weatherby.genetics.utah.edu/phevor2/index.html). The output of Phevor contains a list of ranked genes, enabling direct comparison with AMELIE. The Phevor gene rank was calculated as the number of candidate causative genes ranked before the causative gene plus 1.

#### Comparison to Phenomizer

The output of Phenomizer^30^ for each of the 215 patients was obtained through Phenomizer's website (http://compbio.charite.de/phenomizer/). The output of Phenomizer consists of a list of ranked diseases along with the set of genes known to be associated with each disease. In contrast, AMELIE's output consists of a list of genes along with the articles that explain why mutations in this gene could be causing the patient's phenotype.

To compare the output of Phenomizer with AMELIE's output, the rank of any gene from Phenomizer's output was calculated as follows: (1) Take the patient's disease as the highest Phenomizer-ranked disease associated with the causative gene. (2) Take the set of all genes associated with a disease at higher Phenomizer rank than the patient's disease. (3) Let *x* be the number of unique genes found in step (2). (4) The rank of the causative gene equals *x* + 1.

### Performance of AMELIE is unchanged when training only on 5-year-old data

The publication year of an article was taken from the publication date of the original article, which is saved in PubMed. The relevant document classifier and the topical gene classifier were trained on PubMed titles and abstracts from 2011 or earlier. Articles for which the publication date was not deposited in PubMed were omitted from the training data. The training data for the relevant document classifier consisted of 54,537 negative examples and 40,153 positive examples. This is 22% fewer positive training data points compared to the full training set, which contained 51,637 positive examples. The training data for the topical gene classifier consisted of 21,634 positive examples and 263,780 negative examples. This is 29% fewer positive training data points compared to the full training set, which contained 30,291 positive training examples.

### AMELIE holds over 3 times as many gene-phenotype relationships as HPO-A

HGNC gene symbols in the HPO build 115 were converted to gene Ensembl IDs for comparison with AMELIE. The number of gene-phenotypic abnormality links in HPO build 115 is 103,617. Of those, 54,371 were also in the canonicalized set (defined above) of gene-phenotype relationships extracted by AMELIE. Of the remaining 49,246 links, 50 random gene-phenotype links were selected. 33 (66%) out of those were supported by the scientific literature about Mendelian diseases and/or OMIM disease entries for Mendelian diseases. 2 of 50 (4%) were phenotypes linked through cancer, which AMELIE does not attempt to extract. We could not find support for 30% of gene-phenotype links in HPO build 115. Thus, we estimate the number of true gene-phenotype links for Mendelian diseases that AMELIE is missing to be 66% of 49,246, or 32,502. The number of gene-phenotype links extracted by AMELIE that are not in the canonicalized version of HPO build 115 is 588,595. By sampling 50 random gene-phenotype associations out of those, 18/50 (36%) were determined to be correct extractions. The estimated number of true extractions in AMELIE that are not in the canonicalized version of HPO build 115 is therefore 588,595 * 36% = 211,894.

